# Quantifying the Structural and Energetic Consequences of EXOSC3 S1 Domain Variants from a Comparative Assessment of λ-Dynamics with Two Charge-Changing Perturbation Strategies

**DOI:** 10.1101/2025.06.30.662390

**Authors:** Monica P. Barron, H.R. Sagara Wijeratne, Avery M. Runnebohm, Katelyn M. Caric, Amber L. Mosley, Jonah Z. Vilseck

**Author notes:** **CORRESPONDING AUTHOR Jonah Z. Vilseck (Email:**).

## Abstract

Charge-changing perturbations are notoriously difficult to investigate with alchemical free energy calculations. The routine use of periodic boundary conditions and electrostatic approximations, such as particle-mesh Ewald (PME), may produce finite-size effect errors that become non-negligible as a perturbation changes a simulation cell’s net charge away from zero. Two prevalent strategies exist to correct for these errors: the analytic correction (AC) and co-alchemical ion (CI) methods. Both correction schemes have been found to produce comparable relative free energy results for small molecule perturbations, but these methods have not been compared using λ-dynamics (λD) free energy calculations or for protein side chain mutations. Recently, we investigated relative folding and binding free energies (ΔΔ*G*s) of a series of EXOSC3 variants involved in a rare neurodegenerative disorder, including D132A, G135R, and G191D charge-change perturbations, with a simplified AC scheme in λD. In this study, these perturbations are reevaluated with the CI scheme for comparison with AC to identify the best correction strategy for λD. The collected AC- and CI-corrected ΔΔ*G*s show excellent agreement with a mean unsigned error of 0.4 kcal/mol. However, reduced sampling proficiency and increased difficulties of evaluating multisite perturbations with the CI method suggest that a simplified AC approach may be more generalizable for future λD calculations. Previously, the use of the CI approach with λD has been limited due to a lack of infrastructure available to users to simplify its more involved setup procedure. This study introduces an automated workflow for implementation of the CI approach with λD, laying the foundation for future comparisons between charge-change correction schemes. These studies facilitated analysis of the λD trajectories to identify structural changes within EXOSC3 and the RNA exosome complex that clearly rationalize the calculated ΔΔ*G*s for the D132A, G135R, G191C, and G191D EXOSC3 variants, providing insight into potential disease-causing mechanisms of EXOSC3 modifications.

## INTRODUCTION

Proteins, with their abundant and diverse functionality, play fundamental roles as essential molecular machines in practically all biological processes. [1] Furthermore, studies suggest that around 80% of proteins perform their primary functions by interacting with other proteins. [2] Hence, it is not surprising that altered protein-protein interactions (PPIs) can play devastating roles in various diseases, including neurodegeneration, cardiovascular diseases, metabolic disorders, and cancer. [3–7] PPIs are especially sensitive to single amino acid mutations involved in interfacial contacts between binding partners and can lead to higher likelihoods of disease outcomes. [8] These perturbations can disrupt the binding affinity of one protein for another or the folded stability of one of the interacting partners, thereby lowering the overall frequency or abundance of the native interaction and function of interest. [9]

One such example is found in the RNA exosome. The RNA exosome is a multi-protein complex that is highly conserved and is essential for RNA processing and degradation. The basic exosome complex scaffold is made up of a hexameric barrel ring (subunits named EXOSC4-9) and a trimeric cap ring (EXOSC1-3) (Fig. 1A). From a structural perspective, the exosome core functions as a scaffolding complex, depending on interacting protein partners for enzymatic function. For example, EXOSC10 and DIS3 associate with the cap or base of the exosome, respectively, to perform exo- and endoribonuclease catalytic degradation of RNAs. [10,11–15] Two additional cofactors, helicases SKI2 and MTR4, associate to unwind incoming RNA. [16–21] And the MPP6 exosome cofactor helps tether MTR4 to the exosome by binding EXOSC3 and MTR4. Multiple auxiliary complexes further regulate which substrates are targeted to the RNA exosome complex. [22–24] Amino acid changes in exosome subunits have been associated with tissue-specific diseases, depending on which subunit or cofactor is altered. [25–28] EXOSC3 variants were the first to be identified and associated with disease, specifically pontocerebellar hypoplasia type 1B (PCH1B), a rare neurodegenerative disorder that affects the cerebellum and pons regions of the brain. [25] Pathogenic EXOSC3 variants account for an estimated 50% of known PCH1B cases. [29] Common clinically-observed EXOSC3 variants include G31A and D132A, [25,30–33] with many more variants identified via genomic sequencing, many whose clinical significance is uncertain. [34] Since the discovery of the first PCH1B-linked EXOSC3 variants in 2012, studies have investigated the impact of these variants beyond their clinical phenotype at the organ or tissue level. [35–37] Since many of the first identified variations were found along protein-protein interfaces in the crystal structures of human and yeast exosome complexes, it has been suggested that these EXOSC3 variations disrupted interactions between EXOSC3 and other exosome core subunits. [25,36,37] Two recent multi-omics analyses characterized the molecular and cellular effects of five variations in the S1 domain of EXOSC3, specifically D132A, G135R, A139P, and G191C/D EXOSC3 variants, using RNA sequencing and global proteomics. [38,39] Protein destabilization and significant protein abundance changes were observed in multiple subunits of the RNA exosome complex for each variant, and a high degree of RNA exosome dysregulation was observed via large numbers of changing RNA transcripts. Molecular dynamics simulations were also performed to characterize the molecular mechanisms underlying these variants, revealing specific interactions lost for each variant and providing a rationale for the experimentally observed changes in stability and abundance of EXOSC3 and other proteins associated with the RNA exosome complex. [38,39] This work adds to these molecular characterization studies by more thoroughly examining the thermodynamic consequences of EXOSC3 modification.

**Figure 1.**
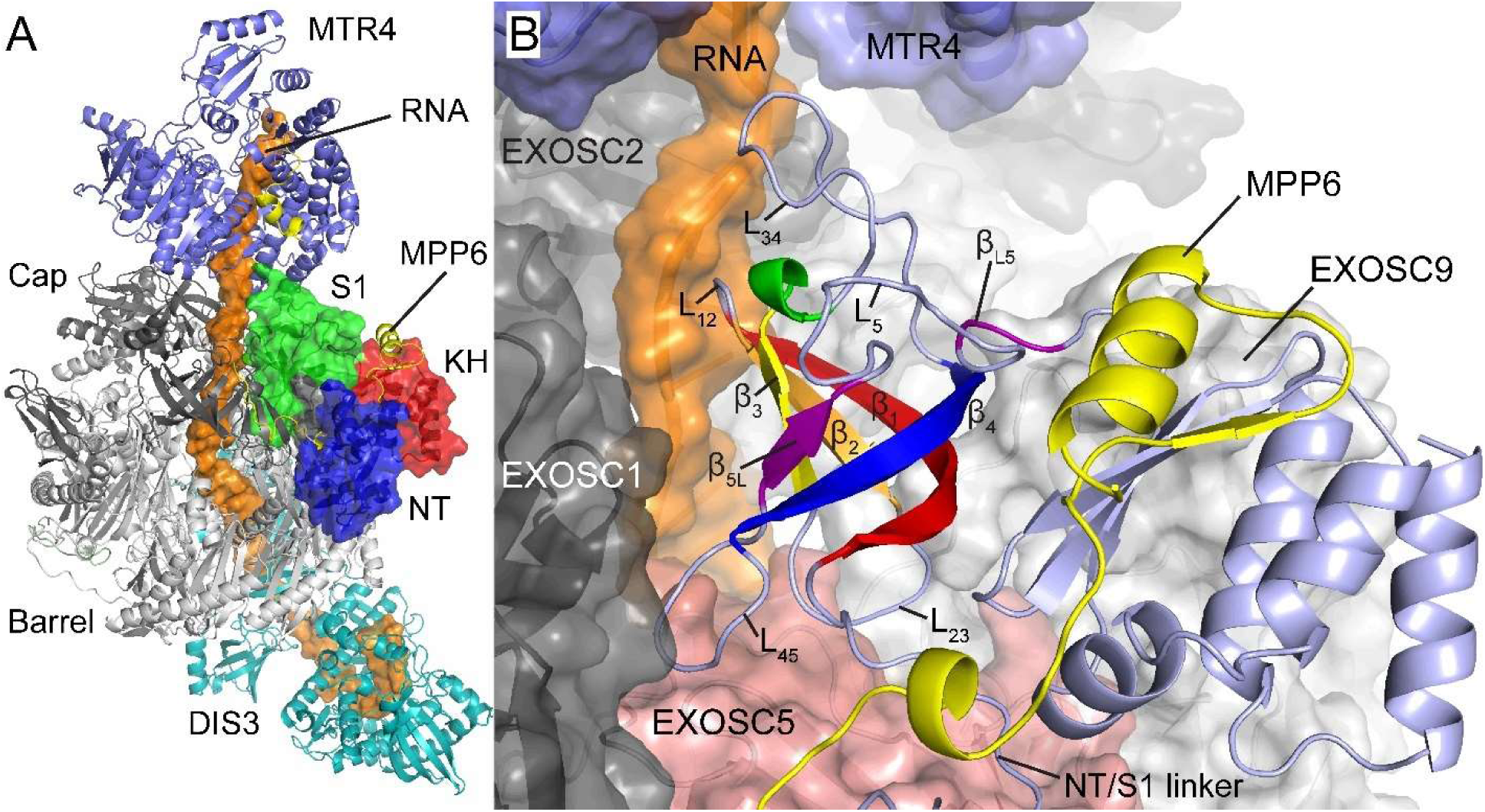
**EXOSC3 structure within the nuclear RNA exosome complex**. (A) A cryo-EM structure (PDBID: 6D6Q) of the human nuclear exosome complex in complex with DIS3, MTR4, and MPP6 cofactors. An RNA strand (orange; surface representation) threads through the exosome channel. The EXOSC3 subunit (surface representation) is colored by domain: NT (blue), S1 (green), and KH (red). All remaining subunits (cartoon representation) are colored as follows: exosome barrel EXOSC4-9 (light gray), exosome cap EXOSC1 and EXOSC2 (dark gray), MTR4 (slate blue), MPP6 (yellow), DIS3 (teal). (B) EXOSC3’s S1 domain in the RNA exosome complex. EXOSC3 is shown as a light blue ribbon. The S1 domain contains a five β-strand barrel labeled β_1_-β_5_ and sequentially colored red, orange, yellow, blue, and purple. A 3_10_ helix (green) follows β_3_. Loops between β-strands are labeled with the strands numbers that flank the loop (i.e. the loop between β_1_ and β_2_ is labeled L_12_). Strand β_5_ is divided into two halves, labeled β_5L_ and β_L5_, respectively, by loop L_5_. Loops L_12_ and L_34_ interact with the RNA in the exosome channel. L_23_ binds the neighboring exosome subunit EXOSC5 (salmon). L_45_ binds both EXOSC5 and the cap subunit EXOSC1. L_5_ interacts with MPP6 and L_34_.

The accurate prediction of the effects of amino acid changes on protein folding or protein-protein binding affinity is crucial for probing the thermodynamic properties of PPIs and understanding how mutations lead to disease. Alchemical free energy calculations can compute free energy differences between native and variant states of a protein sequence, as one sequence is alchemically transformed into another, and, thus, are very useful computational tools to assess thermodynamic properties of biological systems. When performed within a thermodynamic cycle, free energy calculations can compute both relative folding free energies of a monomeric protein, such as EXOSC3 itself, or relative binding free energies of one protein to another, such as EXOSC3 complexed within the larger exosome complex (Fig. 2). Though many free energy methods could be used, λ-dynamics (λD) was chosen for the present work owing to our prior successes using it to study mutational effects in 20S proteasome and insulin-insulin receptor complexes. [40,41] In contrast to other methods, like free energy perturbation (FEP) or thermodynamic integration (TI), λD uses a continuous coupling variable, called λ, to dynamically scale interaction energies between two or more physical end states of a chemical system, such as between multiple amino acid transformations on a given residue. Fortuitously, several transformations can be investigated simultaneously with λD, improving calculation efficiency. Analogous to an experimental competitive binding assay, λ variables in λD dynamically change in conjunction with the forces of a molecular dynamics simulation using extended Lagrangian methods. This enables λD to sample the most favorable perturbations more frequently. [42–44] Free energy landscape flattening is also routinely performed to enable all perturbations to be sampled equally, providing greater comparative mechanistic and structural details between all sampled variants. [45–47] Finally, enhanced dihedral sampling is possible with λD by scaling alchemical side chain dihedral angles by λ. As λ scales to 0, rotational barriers shrink facilitating the rapid sampling of alternative rotameric states without requiring additional enhanced sampling algorithms. [48,49] These collective features make λD a very promising technique for studying the thermodynamic properties of proteins and PPIs.

**Figure 2.**
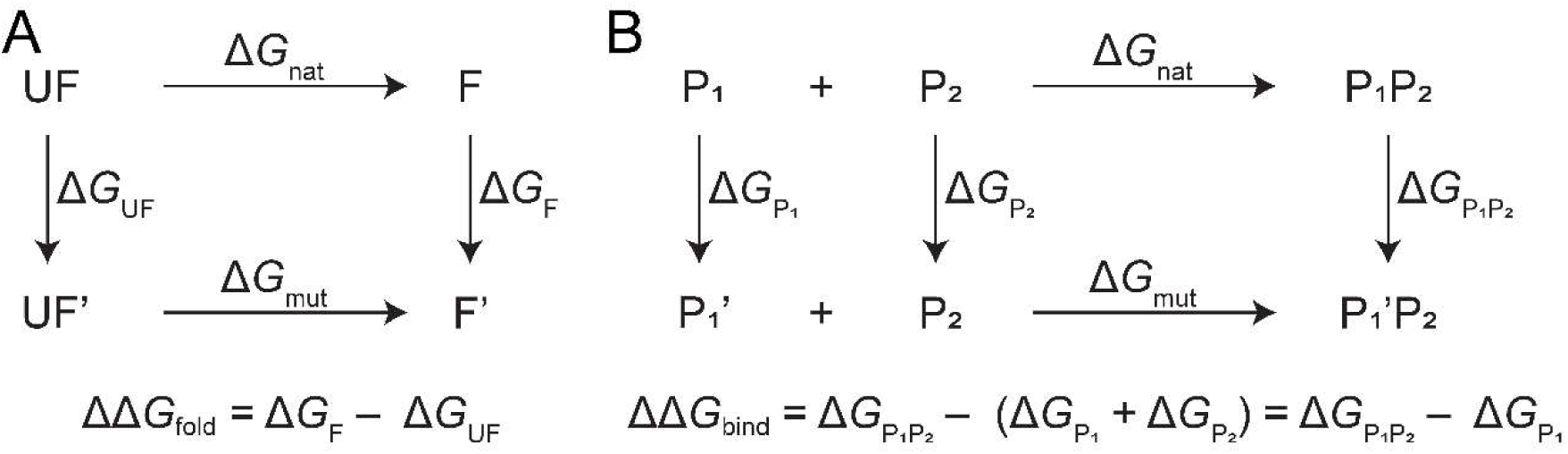
Thermodynamic cycles for calculating relative free energy differences. (A) Relative changes in folding free energies (ΔΔ*G*_fold_) can be calculated by perturbing unfolded (UF) and folded (F) proteins with a native sequence into variant states (UF’ and F’, respectively). (B) Relative changes in binding free energies (ΔΔ*G*_bind_) can be calculated by perturbing one protein (P_1_) into a variant state (P_1_’) in unbound and bound states. P_2_ represents a protein binding partner. Δ*G*_nat_ and Δ*G*_mut_ represent the physical folding or binding processes for native and variant proteins, respectively. ΔΔ*G*_fold_ and ΔΔ*G*_bind_ are computed as a difference of the two alchemical transformations, which equals the natural Δ*G*_nat_ − Δ*G*_mut_ difference for both processes.

In this work, we investigate the binding and folding free energies of D132A, G135R, and G191C/D EXOSC3 variants with λD. However, because three of these transformations involve a perturbation to or from a conventionally ionized residue (Asp or Arg) to a neutral one (Ala, Gly, or Cys), special considerations are required. Perturbations that alter the net charge of a chemical system present a notoriously difficult challenge for alchemical free energy methods. [50–55] To begin with, charged amino acids may have very different interactions with surrounding protein and solvent molecules that are difficult to converge within conventional simulations times (≤ 25 ns per simulation). [50,51] Beyond slow-to-converge structural changes, however, an additional challenge arises from conventional use of a finite simulation box size and periodic boundary conditions in molecular dynamics simulations to represent bulk solvent systems with reduced computational costs, as well as current approaches for treating electrostatic interactions within and between simulation box images, including the particle mesh Ewald lattice sums method for modeling long-ranged electrostatic interactions. [52–57] Perturbing between molecules with different net charges can lead to non-negligible interaction energies between molecules in adjacent simulation cells, disruption of solvent structure in the central cell, undersolvation of alchemically perturbed molecules, and systematic deviations in electrostatic potential energies of charged solutes. [52,53,56,57] Together, these artifacts are collectively referred to as finite-size effects and their magnitude may depend on the size and shape of the system, the treatment of electrostatics, the treatment of simulation cell boundaries, and the ionic strength of the solution. This has been described in detail in the literature. [50,52–54,56,57]

Two dominant strategies exist for correcting finite-size effects accompanying charge-changing perturbations in alchemical free energy calculations: an analytic correction (AC) and a co-alchemical ion (CI) approach. One of the most well-known analytic corrections was proposed by Rocklin *et al.* in 2013 and decomposes finite-size effect errors into periodicity-induced net-charge interactions and undersolvation effects, discrete solvent effects, and residual integrated potential effects. [52] These system-specific corrections factors are computed and applied *post hoc* to an already completed free energy calculation. [50,52–54,57] Rocklin *et al.* demonstrated that the finite-size errors in the charging free energy of a non-neutral ligand are dominated by the discrete solvent effects. [52] As a result, a common simplification of the Rocklin correction is to compute the discrete solvent effects correction only, removing the need to run an expensive Poisson-Boltzmann calculation. [45,58,59] In the co-alchemical ion (CI) approach, charge-changing perturbations are counterbalanced by simultaneously introducing an opposing charge change within the solvent (i.e. perturbing a water molecule to a chloride ion to offset an Asp to Ala variation). [50,51,60] As a mechanical strategy to maintaining a constant system net charge, this approach requires additional changes to the setup and performance of an alchemical free energy calculation but does not require additional post hoc correction terms to be added to the final free energy results. Chen *et al*. found that AC and CI approaches gave comparable accuracies and advocated that the CI approach was more generalizable to arbitrary new systems of interest. [50]

Although, AC and CI strategies have been evaluated with traditional free energy calculations, including FEP and TI methods, a comparison of the two correction schemes had not yet been performed within the λD framework. Because λD fundamentally differs from FEP or TI via its use of a dynamic, continuous λ variable, it was not readily apparent if the CI approach would work better for λD than the AC scheme, as reported for other methods. [50,55] We investigated relative changes in folding free energies (ΔΔ*G*_fold_) and binding free energies (ΔΔ*G*_bind_) of a series of variations within the S1 Domain of EXOSC3, specifically D132A, G135R, G191C, and G191D, with the simplified Rocklin analytic correction. [38,39] In this manuscript, we introduce an implementation of the CI approach for λD and repeat ΔΔ*G*_fold_ and ΔΔ*G*_bind_ λD calculations using this CI correction strategy. We compare the free energy results obtained using the CI correction strategy to those obtained using the simplified AC correction strategy. The λD setup, observed performance, and structural results are each compared between the two methods. Sampling and convergence challenges accompanying the analyzed modifications are discussed along with improvement strategies. Thus, this work establishes a side-by-side comparative assessment of the CI and simplified AC schemes for use in λD calculations, applied to an analysis of EXOSC3 S1 domain missense variants involved in PCH1B. [45,58,61] In this comparison, we aim to provide insight into the setup, sampling, and analysis of these two correction schemes, and provide guidance on the selection of an appropriate strategy for future λD experiments.

## RESULTS

### λD implementation of the co-alchemical ion correction strategy

As described above, the CI scheme for maintaining net system charge neutrality and negating finite-size effect errors involves coupling a primary alchemical transformation with a charge change, in this case an amino acid side chain, with a secondary transformation with an opposite charge change in the solvent environment. [50,51] Three different CI variants have been reported in the literature. In general, a charged ion in the solvent can be neutralized by mutating it into a null dummy particle with no charge or Lennard–Jones parameters, a Lennard–Jones particle matching the original ion’s identify with no charge, or a water molecule. [50] As the last approach has been suggested to be the most robust, the co-alchemical ion-to-water variant of the CI method was selected for implementation in λD. [50]

An automated CI protocol was created to facilitate use of this correction scheme with λD without requiring manual adjustments to basic λD scripts. For most protein side chain or ligand functional group perturbations, typical charge-change perturbations comprise a change in charge (Δq) of only ± 1 e^-^. For protein perturbations between native amino acids, however, perturbations between positively and negatively charged residues would result in a Δq of ± 2 e^-^. Thus, in our automated protocol, for instances of Δq = ± 1 e^-^, charged substituents are each coupled to a water molecule and neutral substituents are each coupled to an ion of the same charge as the charged substituent(s). This precludes any challenges that might arise when two charged species are both introduced or both removed simultaneously. In instances of Δq = ± 2 e^-^, charged substituents are each coupled to an ion of the opposite charge and neutral substituents are each coupled to a water molecule. This allows the charge change to be counteracted by perturbations of a single solvent molecule. These assignments depend upon the set of perturbations to be sampled within a single λD simulation and are made independently of the charge or identify of the native substituent. In all cases, the ion or water molecule to be alchemically coupled is randomly selected from the solvent box as an ion or water molecule that is the farthest away from the center of the box but within a user-defined distance (default 2 Å) from the edge of the simulation box. Further practical details of CI implementation in λD scripts are presented in the Discussion and Methods sections.

### Excellent thermodynamic agreement was obtained between the two charge-change correction strategies

CI-corrected λD simulations of EXOSC3 charge-change perturbations were set up in a similar manner as associated AC-corrected λD simulations of the same perturbations. [38,39] The AC-corrected simulations used the simplified approach and corrected for discrete solvent effects only. λD production simulations were run for exosome-bound, unbound, and unfolded EXOSC3 systems in the CHARMM molecular modeling software package. [62–64] Separate λD calculations were run to model the perturbations of each EXOSC3 residue modified (D132, G135, and G191). To offset the difficulty of the G135R transition, two additional perturbation states were included in the G135 perturbation set, α-aminobutyric acid (G135Abu) and norleucine (G135Nle). [39] Similarly, a perturbation to alanine (G191A) was included in the G191 perturbation set to offset the transition from Gly to Cys or Asp. [39] These offsets provide gradual volumetric steps for λD to naturally transition from a small, flexible Gly residue into larger, charged side chains and can aid protein response and relaxation along the transformation pathway. Production λD simulations were run with 4-5 replicas for a minimum of 100 ns each, or until converged free energies were obtained (up to a maximum simulation length of 200 ns per replica). Plots of computed free energies with respect to time over the last 50 ns of each simulation confirmed that the free energy results were converged (Figs. S1-S3). Additional computational details are reported in the Methods section.

As shown in Figure 3 and Table S1, the thermodynamic results displayed excellent agreement between the AC and CI charge-change correction schemes, with a mean unsigned error (MUE) of 0.40 kcal/mol over all fourteen perturbations and a Pearson R correlation of 0.98. Nine perturbations showed very good agreement within ± 0.5 kcal/mol (dark gray band), while the remaining perturbations were within ± 1.0 kcal/mol (light gray band). The average uncertainty for the data set was 0.25 kcal/mol, with some uncertainties ≤ 0.1 kcal/mol and a maximum uncertainty of 1.0 kcal/mol for G135, owing to sampling difficulties encountered for this specific perturbation. The free energy results suggest that the D132 and G191 variants affect EXOSC3 folding and stability more significantly than binding, whereas the opposite is true for G135. This is discussed in detail with additional structural analyses below. The high agreement between AC and CI λD results in this work match conclusions reported from other comparisons with different alchemical free energy methods, [50,55] and confirms that both strategies work well for negating finite-size effect errors to yield converged, reproducible free energy differences. With this high level of agreement in hand, effects on λD bias determination, sampling proficiency, and convergence were analyzed to determine if one approach was better suited for λD than the other.

**Figure 3:**
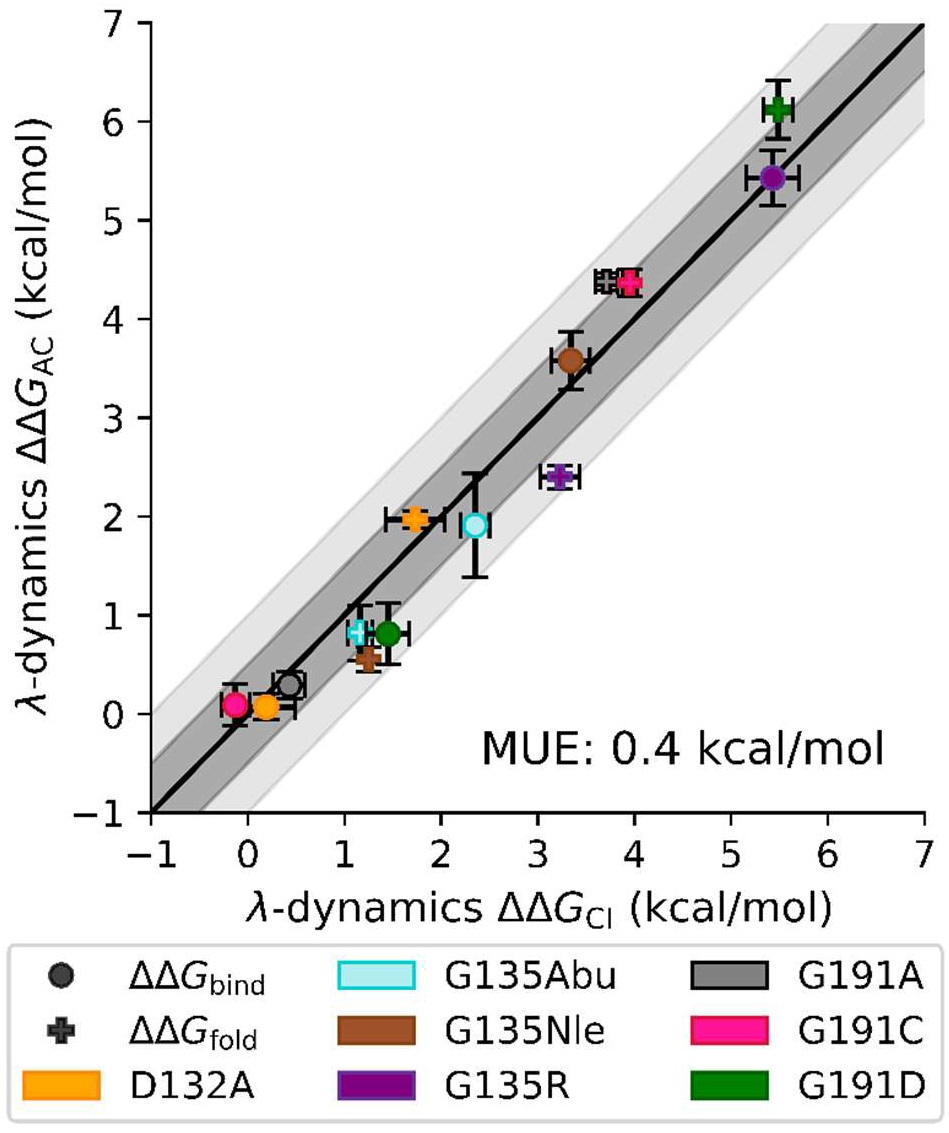
Comparison between CI and AC charge-correction protocols of λD computed EXOSC3 folding and exosome-binding free energies. Relative binding affinities (ΔΔ*G*_bind_, ●) and folding free energies (ΔΔ*G*_fold_, ✚) were computed with λD using either a co-alchemical ion (CI, X-axis) or analytic correction (AC, Y-axis) scheme to counter finite-size effect errors accompanying charge-changing D132A, G135R, and G191D perturbations. Intermediate and charge-preserving perturbations, including G135Abu, G135Nle, G191A, and G191C, are also shown. The solid black line indicates perfect agreement between the two methods (y=x). The dark gray and light gray bands represent ΔΔ*G* errors within ± 0.5 kcal/mol and ± 1.0 kcal/mol, respectively.

### The charge-change correction strategy does not influence the length of pre-production bias determination sampling

The impact on λD bias determination was evaluated first. As is routine for λD simulations, the Adaptive Landscape Flattening (ALF) algorithm was used to identify appropriate biasing potentials for dynamic sampling of multiple end states (see Methods). [46,47] To investigate if the charge-change correction strategy impacted the amount of pre-production sampling required to optimize λ biases, the total amount of pre-production sampling for the two correction schemes was compared (Table S2). On average, CI simulations used 420 ns of sampling, while AC simulations required only 334.5 ns. The amount of ALF pre-production time for modeling individual variants ranged from ca. 170 - 800 ns. However, this significant variance in sampling appeared to be more related to the impact of the modification on the surrounding protein structure than the charge-change correction strategy. For example, the G135 and G191 variants both included small-to-large perturbations accompanied by loss of backbone flexibility unique to Gly. In the unbound, folded EXOSC3 subunit and the RNA exosome complex, these modifications led to large structural rearrangements (see below) that affected protein packing and were difficult to converge. Protein relaxation and local conformational changes to accommodate these perturbations were expected to require longer time scales of sampling. A more representative comparison of pre-production simulation time difference may be obtained by limiting the comparison to the D132 perturbations and unfolded pentapeptide simulations of the G135 and G191 perturbations, since slow-to-converge structural changes were not observed in these simulations. From this subset of simulations, the required amount of pre-production ALF sampling ranged from 170 - 300 ns, with very similar average amounts of sampling for the CI simulations (231.8 ns) compared to the AC simulations (250.8 ns). This implies that the choice of charge-change correction strategy does not greatly impact the amount of ALF sampling necessary to optimize λ biases, but rather the identify and packing location of perturbations explored are more important factors.

#### The CI strategy modestly decreased λD transitions and substituents sampling statistics

To accurately calculate free energy differences between two or more end states with λD, it is necessary to sample each end state many times and frequently transition between all substituents. [46] Free energy differences can then be calculated as a ratio of the amount of sampling of one end state with respect to a reference state (equation 1),

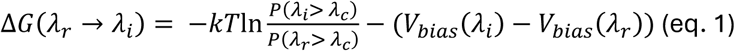

where 𝜆_i_ is the coupling parameter associated with perturbation end state 𝑖, 𝜆_r_ is the coupling parameter associated with the reference state, 𝜆_c_ is the cutoff value used to define perturbation end states (typically a λ_c_ > 0.99 is used), and 𝑉_bias_(𝜆) is the biasing potential for a given λ state (λ_i_ or λ_r_). To aid in meeting this sampling requirement for λD, free energy barriers in λ space are flattened by system-specific biases optimized by ALF, which should enable all end states to be sampled nearly equally during a λD simulation. In practice, however, perfectly equal sampling is rarely observed due to system variation and visiting new conformational states not previously observed in pre-production simulations. As a rule of thumb, to gauge if λD sampling will yield converged free energy results, we typically aim to maintain a sampling ratio between the most and least sampled states that is within a factor of 10. To determine the effect of CI vs AC correction schemes on λD sampling proficiency and convergence, the number of transitions between end states and the percent each perturbation state was sampled were tabulated and compared (Table S3). A larger average number of transitions indicates improved sampling performance. Sampling percentages should be equivalent between perturbations sampled within a single λD calculation, within a factor of 10, and larger percentages are preferred because they represent more frequent end state sampling over alchemical intermediates. From Table S3, even sampling of all substituents was achieved across all λD simulations with the exception of both the CI and AC simulations of the exosome-bound EXOSC3 G135 variants, where suboptimal sampling was observed for the G135Abu and G135Nle intermediate states. However, the CI and AC ΔΔ*G*_bind_ values of the clinically relevant G135R variant were unaffected since sufficient sampling was obtained for both the G135R variant and the reference G135 end states. A clear relationship was observed between the number of transitions and the EXOSC3 model (exosome-bound, unbound, and unfolded), with decreasing model size and complexity accompanied by an increased number of transitions. This relationship maps the increased difficulty of introducing an amino acid change within a protein complex relative to the less constrained protein monomer or the flexible pentapeptide. As expected, an increase in the average number of transitions was accompanied by a modest decrease in the average sampling percentages, indicating that the system was less likely to become trapped in any one end state. Compared to the AC simulations, the CI simulations displayed decreased average numbers of transitions, ca. 385 transitions per 100 ns, compared to the ca. 590 transitions per 100 ns observed with AC simulations. Slight decreases were also observed in the average sampling percentages for CI perturbations. Since the CI approach couples two alchemical perturbations together, some additional sampling difficulties are expected; however, it did not appear to affect the ultimate ΔΔ*G*_bind_ or ΔΔ*G*_fold_ results. This difficulty is likely to be compounded in larger, multisite studies, where charge-change perturbations may be performed at two or more sites along a protein sequence. This trend could make the AC approach a more appealing choice for these systems, since multisite systems can sometimes struggle to achieve desired sampling ratios and transition rates, due to the increased combinatorial alchemical space explored. [45]

### Structural analyses of λD trajectories rationalize the computed thermodynamic changes accompanying EXOSC3 variation

As described earlier, the EXOSC3 protein is a cap subunit of the RNA exosome complex and contains three domains: an N-terminal domain (NT), an RNA-interacting S1 domain, and a KH domain (Fig. 1). The perturbed residues studied herein, D132, G135, and G191, all lie within the EXOSC3 S1 domain, which is made up of a β-barrel containing five β-strands, labeled β_1_-β_5_ (Fig. 1B). While the D132 and G135 perturbation sites lie within L_23_, the loop between the β_2_ and β_3_ strands, the G191 perturbation site is found within β_L5_, a subsection of the β_5_ strand. Structural consequences of the EXOSC3 S1 domain variants were extracted via analysis of CI and AC λD trajectories to investigate the origin of the free energy results reported above, with each variant analyzed independently.

#### EXOSC3 D132A missense variant

Investigation of the D132 λD trajectory frames revealed that a consistent hydrogen bond is formed between the D132 carboxylate side chain and the G134 backbone nitrogen that helps stabilize the native configuration of the L_23_ backbone conformation (Fig. 4A). Loss of this hydrogen bond in the D132A variant is associated with increased flexibility and loss of the native L_23_ conformation (Fig. S4A). [38] Alternative conformations of L_23_ (Fig. 4B) cause two additional stabilizing hydrogen bonds between the NT/S1 linker (the linker between the EXOSC3 NT and S1 domains) and L_23_ to break, including G134 O-Y109 N and R108 guanidine to L_23_ Gly134/135 backbone oxygen interactions (Fig. S4B). This change in configuration of L_23_ and the resulting disruption of stabilizing hydrogen bonds between the NT/S1 linker and the L_23_ backbone is accompanied by a ΔΔ*G*_fold_ of ca. 2 kcal/mol for D132A EXOSC3. Because the ΔΔ*G*_bind_ was near 0.1 kcal/mol, these results suggest D132A primarily destabilizes EXOSC3’s monomeric structure, but once folded, minimally impacts its binding affinity to the exosome complex.

**Figure 4.**
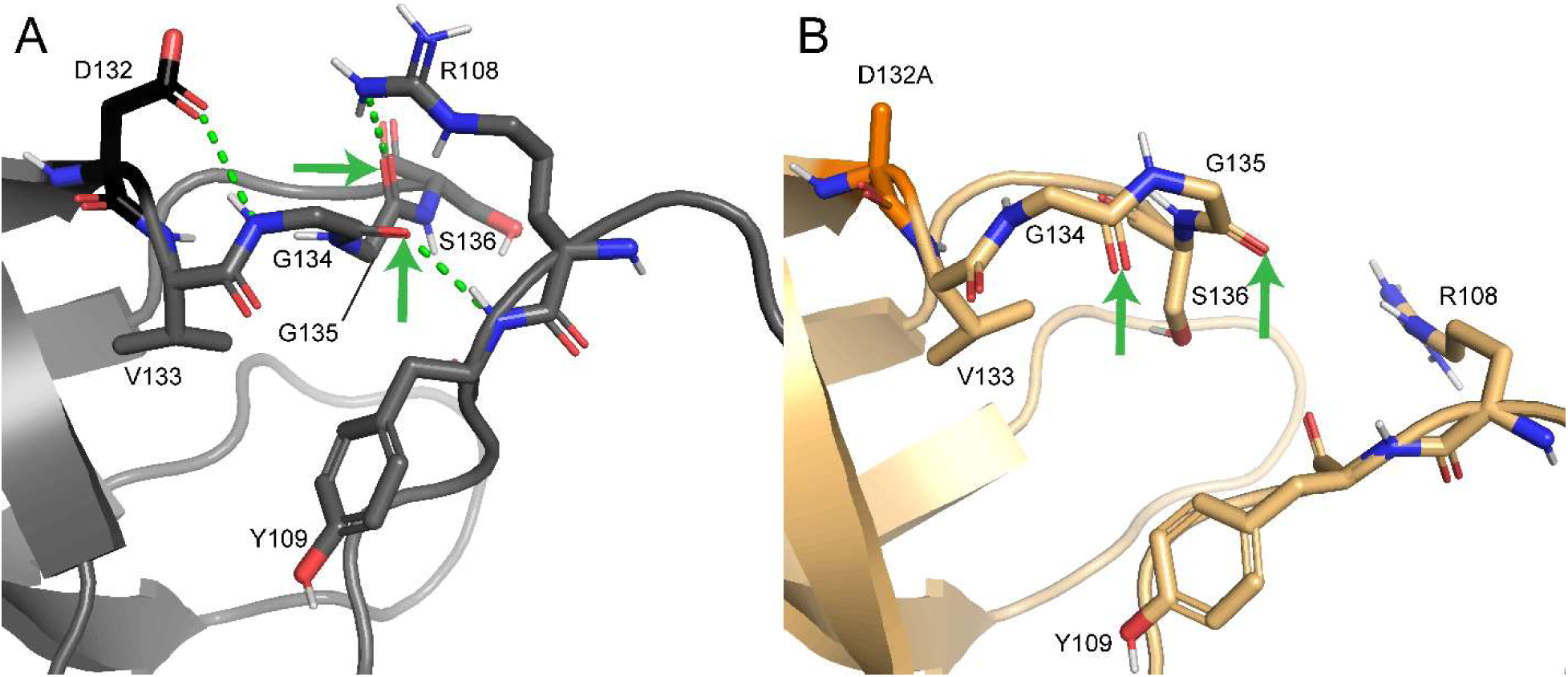
Structural changes introduced by the EXOSC3 D132A variant. (A-B) Sample frames of unbound D132 EXOSC3 with the native L_23_ configuration (A) and unbound D132A EXOSC3 with an alternative L_23_ configuration (B). Green arrows highlight orientational changes in the G134 and G135 backbone in the alternative L_23_ configuration. Hydrogen bonds observed in native D132, but which are broken in D132A, are shown as green dashed lines.

#### EXOSC3 G135R missense variant

The G135 residue is positioned in the center of L_23_ and packs against NT/S1 linker residues R108 and Y109. In the RNA exosome complex, both L_23_ and the NT/S1 linker pack tightly against EXOSC5, a neighboring exosome barrel subunit. Representative structures of both exosome-bound and unbound EXOSC3 were collected from the G135 and G135R frames of the CI-corrected λD simulations (Figs. 5 and S5), which also match structures observed in the AC-corrected simulations. In exosome-bound G135 EXOSC3, tight packing is observed between L_23_, the NT/S1 linker, and EXOSC5 (Fig. 5A). The G135R variant, however, directly clashes with the NT/S1 linker and disrupts many interactions with L_23_ and EXOSC5, as seen by visible opening between adjacent loops (Fig. 5B). This large structural distortion of EXOSC3 packing leads to a large ΔΔ*G*_bind_ of ca. 5.5 kcal/mol for G135R in the exosome complex. In unbound EXOSC3, similar packing distortions are observed (Fig. S5A), but the magnitude of the ΔΔ*G*_fold_ is reduced to ca. 2-3 kcal/mol, since the structure of EXOSC3 is not constrained by surrounding PPIs.

**Figure 5.**
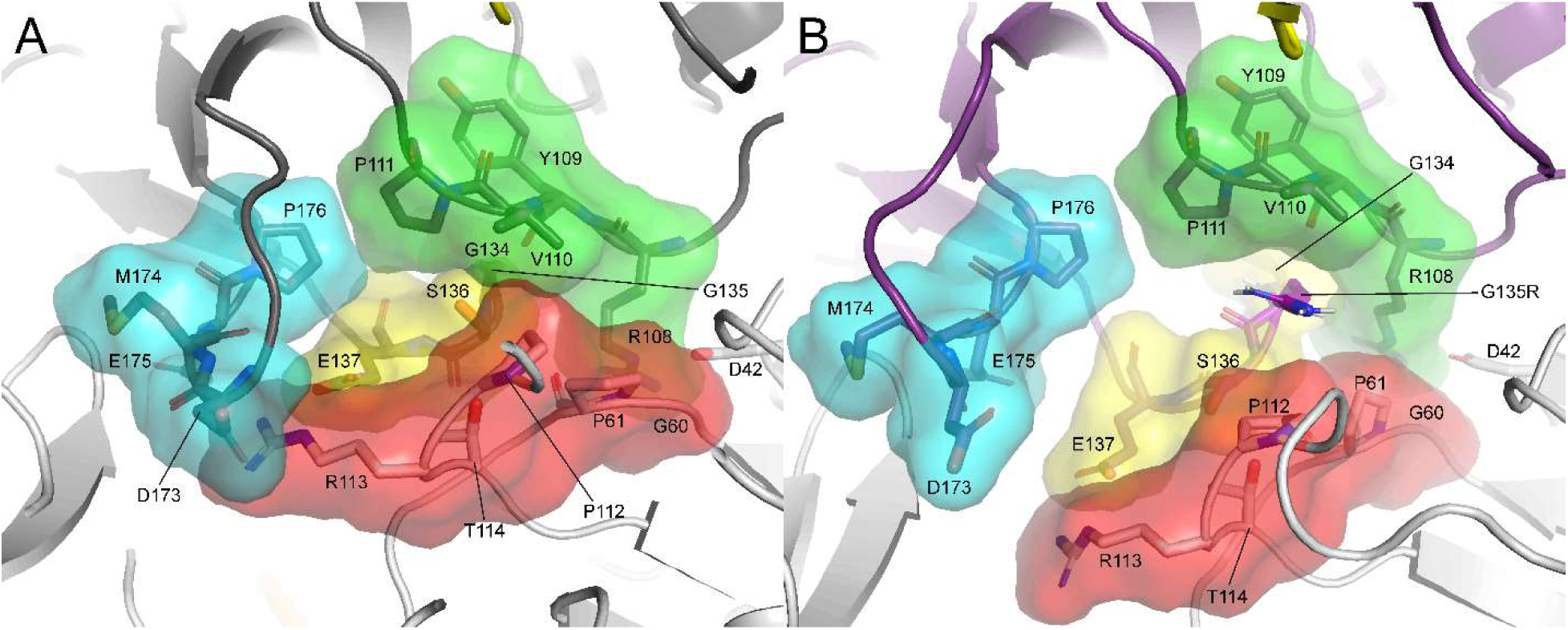
Disruption of packing interactions in exosome complex by EXOSC3 G135R variation. (A) Tight packing between EXOSC3’s NT/S1 linker (green), L_23_ (yellow), L_45_ (cyan), and EXOSC5 loops (red) for native EXOSC3 (dark grey). (B) Disrupted packing in G135R EXOSC3 (purple). Representative frames were collected from CI-corrected λD simulations of exosome-bound EXOSC3.

#### EXOSC3 G191C/D missense variants

The G191 residue sits at the beginning of β_L5_, a three-residue continuation of the β_5_ strand located after L_5_ (Fig. 6A) and forms two of β_L5_’s three hydrogen bonds with β_4_ (Fig. 7A). The G191 hydrogen bonds and the native configuration of the G191 backbone are critical for the structure and behavior of L_5_, [39] especially in unbound EXOSC3, but are unfavorable for non-glycine amino acids. Investigation of the G191 variant λD trajectory frames reveals that G191D (Fig. 6B) and, at a lesser frequency, G191C (Fig. 6C) disrupt all three of these β_L5_-β_4_ hydrogen bonds (Fig. 7B) and cause the G191C/D and V192 residues to detach from a hydrophobic surface that G191 contacts. [39] The loss of β_L5_ structural integrity further alters the structure and behavior of L_5_. Together these changes are accompanied by a ΔΔ*G*_fold_ of ca. 6 kcal/mol for G191D and ca. 4 kcal/mol for G191C (Fig. 3). Relative binding free energies for G191D (ca. 1 kcal/mol) and G191C (ca. 0 kcal/mol) were much lower, suggesting that the structural changes accompanying the G191 perturbations primarily destabilize EXOSC3’s folded monomer with a less significant impact on the stability of the exosome complex.

**Figure 6.**
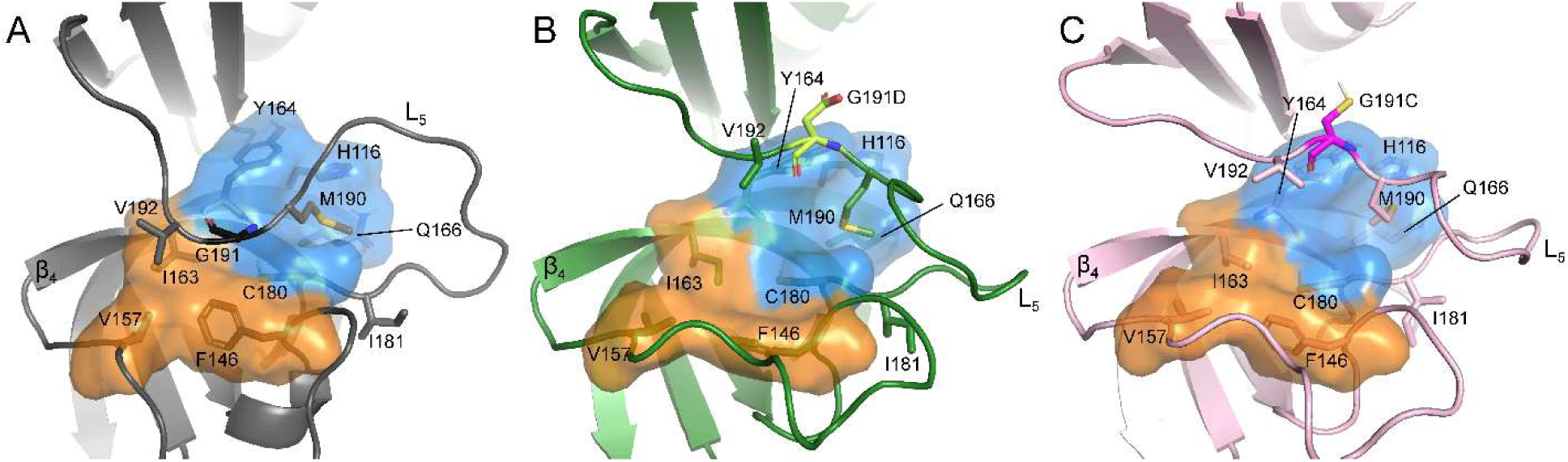
Disruption of the β_L5_ structure by the G191 variants. (A-C) The G191C/D variants disrupt hydrophobic interactions near β_L5_ and lead to a loss of hydrogen bonding near the 191 residue. (A) In native G191, M190 binds within a hydrophobic pocket formed by H116, Y164, Q165, and C180 (blue surface). G191 rests against a hydrophobic surface made up by C180, I163, and frequently F146 (colored with blue and orange surfaces respectively). And V192 interacts with I163, F146, and V157 (orange surface). G191D (B) and G191C (C) variants break contacts between β_L5_ and the colored hydrophobic surfaces and alter the dynamic behavior of L_5_.

**Figure 7.**
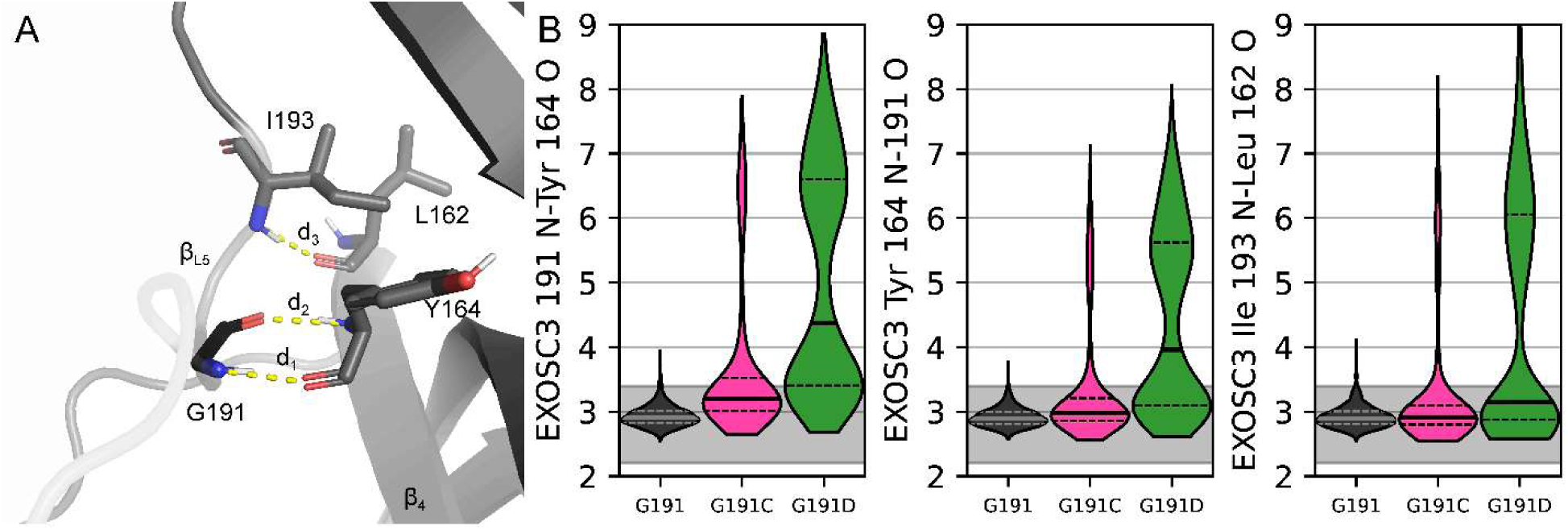
**Disruption of β_L5_-β_4_ hydrogen bonds by the G191 variants**. (A) A representative G191 structure highlights three key hydrogen bonds that connect the β_L5_ and β_4_ strands, labelled d_1_ (191 N-Y164 O), d_2_ (191 O-Y164 N), and d_3_ (I193 N-L162 O). (B) Distributions of hydrogen bonding distances for G191 (black violins), G191C (pink violins), and G191D (green violin) variants collected from CI-corrected λD production frames of unbound EXOSC3. Left panel: 191 N – Y164 O (d_1_, Å). Middle panel: 191 O – Y164 N (d_2_, Å). Right panel: I193 N – L162 O (d_3_, Å). All violin plots are normalized to have the same width. Solid lines on the violin plots represent median values and dashed lines show the interquartile range. The gray band indicates a typical hydrogen bonding distance of 2.2-3.4 Å between heavy atoms.

## DISCUSSION

Although both the CI and AC charge-change correction strategies have been successfully implemented in alchemical free energy simulations for many years now, the simplified AC strategy has been predominantly used to correct for charge change perturbations in λD. Considering that the CI strategy has become a correction strategy of choice for many alchemical free energy methods, such as FEP+, we sought to evaluate its performance with λD compared to the simplified AC approach. However, the relative difficulty of CI simulation setup can quickly increase as multiple perturbations are performed at one or more sites in a λD simulation and within large studies with many different charge-changing perturbations. For example, in this study, the three charge-changing EXOSC3 perturbations (D132A, G135R, and G191D) required nine different series of calculations to be set up with the CI charge-correction scheme to calculate the desired ΔΔ*G*_fold_ and ΔΔ*G*_bind_ properties for each EXOSC3 variant. The expansion of perturbations could be significant when analyzing patient-specific protein dynamics.

In the absence of a scripted, automated setup for CI-corrected λD simulations, a user would have to manually carry out the following steps. First, assess each perturbation site to identify the presence of a charge-change perturbation as well as the direction and magnitude of the charge change involved. Second, select an appropriate solvent perturbation scheme. For example, if the co-alchemical ion-to-water variant of the CI method is chosen, a charge-change perturbation of Δq = + 1 e^-^ performed in a solvent box with NaCl counterions could be neutralized by perturbing either a reference water molecule to a sodium ion or by a reference chloride ion to a water molecule. Similarly, a Δq = + 2 e^-^ perturbation could be countered by perturbing a reference chloride to a sodium ion or by introducing two water-to-sodium or two chloride-to-water perturbations simultaneously.

Next, the reference solvent molecule or ion to perturb needs to be selected from the solvated system. Random selection of the reference molecule is discouraged as it could end up in the vicinity of the protein or ligand near the center of the box, which may introduce additional artifacts into the computed free energy differences. [50] Fourth, for each substituent at the charge-change perturbation site, an appropriate co-alchemical water or ion needs to be generated. Finally, the λD CHARMM input script needs to be modified to pair each alchemical ions or waters with its appropriate primary substituent or side chain partner in the BLOCK module and to restrain the co-alchemical heavy atoms together using the constrained atoms (*CATS*) function in CHARMM. [65] Within our automated CI protocol, each of these steps is fully automated for the user, making the CI correction strategy as easy to implement as the post-simulation simplified AC method, with minimal user intervention or processing.

Challenges arise for the CI approach in systems investigating charge-change perturbations at multiple sites. This limitation is related to challenges in the positioning of alchemical solvent atoms within the system. Ideally, co-alchemical solvent molecules should be positioned outside the normal electrostatic cutoff distance (typically set at 12 Å) relative to all other charge-change perturbation sites, whether those perturbation sites are in the ligand/macromolecule or in the solvent. The ability to separate the charge-change perturbation sites would become difficult should multiple co-alchemical solvent molecules be included in the system, especially in smaller systems. The script currently selects reference solvent molecules or ions from the input structure such that the value 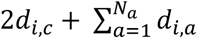 is maximixed, where 𝑑_i,c_ is the distance between the alchemical solvent molecule to be placed and the center of the simulation box and 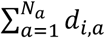 is the sum of the distance between the alchemical solvent molecule to be placed and all previously placed alchemical solvent molecules. However, this operation does not account for co-alchemical ions duplicated across periodic image lattices. Consequently, the current version of this protocol is not recommended for any simulation that requires more than one or two co-alchemical ions to be introduced in a simulation, such as exploring multiple charge-change perturbations on different protein residues simultaneously. This protocol also assumes a cubic solvent box when determining the edge of the water box. If a non-cubic water box is selected, a co-alchemical solvent molecule might be placed at the very edge of the water box, which could potentially lead to artifacts in the free energy calculation if the selected solvent molecule is restrained at the periphery between periodic images. The co-alchemical solvent molecules in this study were allowed to freely move within the boundaries of the water box. Finally, this protocol assumes that the salt buffer is composed of monovalent ions only (i.e. NaCl or KCl). The automation of the CI correction scheme greatly simplifies the performance of λD simulations with charge-changing perturbations by automating time-consuming steps that previously had to be carried out by a user for every λD simulation utilizing the CI correction scheme.

This CI automation scheme also greatly assists comparisons between CI and AC approaches, the two predominant charge-change correction schemes, to evaluate which functions better for λD specifically. Comparing pre-production simulation times for each variant shows the charge-change correction strategy does not significantly affect ALF pre-production sampling (Table S2). Instead, pre-production simulation times depended more on the nature of the perturbations, such as larger amino acid perturbations and accompanying structural changes in the protein complex. In the production λD simulations, the CI method consistently decreased the number of transitions between substituent end states as well as the amount of end state sampling, compared to AC simulations (Table S3). This suggests that sampling was slightly improved with the AC approach than with CI. Nevertheless, high ΔΔ*G* agreement was obtained between the two charge-correction strategies, with a mean-unsigned error of 0.4 kcal/mol across all fourteen thermodynamic results calculated (Fig. 3 and Table S1). Although the CI correction strategy did not significantly alter the required pre-production simulation time or the calculated ΔΔ*G* results, the observed decreases in sampling and transitions could become problematic for simulations sampling a large number of perturbations at a single site or when performing perturbations at two or more sites. In multisite λD, slower transition rates could prolong convergence in free energy estimates. Furthermore, coupling charge-changing perturbations at more than one site quickly increases the complexity of a λD simulation by requiring multiple independent co-alchemical ions to be placed in the system. In contrast, the simplified AC correction scheme enables many charge-changing perturbations to be performed simultaneously, regardless of how many sites are mutating at once. This generalizability for sampling multiple perturbations at one or more than one site naturally fits the sampling capabilities of λD, known to improve free energy calculation efficiency and scalability. Combined with improved sampling characteristics, including sampling ratios and high transitions rates, the AC correction scheme is better suited for most λD simulations, especially screens involving many amino acid side chain substitutions. [42,45] Single site CI λD calculations may continue to play important roles in small, isolated investigations or for confirming the reproducibility of AC λD results.

Beyond performing a comparative assessment of the CI and AC charge-changing perturbation strategies in λD, this study also investigated the structural and energetic consequences of the EXOSC3 D132A, G135R, and G191C/D variants using λD. Structural analyses of the λD simulation trajectories provided a clear rationale for the calculated ΔΔ*G*s for EXOSC3 variants, as presented in the Results section. Furthermore, the accompanying structural changes identified in this study are remarkably similar to those identified by MD simulations of the D132A, G135R, G191C, and G191D variants. [38,39] This reproducibility in structural descriptions demonstrates the power of λD for investigating the structural origin of calculated folding and binding free energies.

In conclusion, to our knowledge, this study represents the first reported comparison of CI and AC charge-change correction strategies using λD free energy calculations. Findings were then applied to the analysis of clinically identified EXOSC3 S1 domain missense variants involved in PCH1B. An automated implementation was developed to assist in CI setup for λD, and CI vs AC performance was measured. The two approaches were found to produce statistically equivalent thermodynamic results with an MUE of 0.4 kcal/mol, suggesting high reproducibility in both correction schemes with λD. Yet the CI perturbation was consistently found to modestly decrease the number of transitions between end states and sampling ratios, suggesting slightly improved sampling was obtained with the AC approach. Considering the expansive sampling capabilities of λD suggests the simplified AC approach may be a more generalizable approach for future λD simulations. However, because this study only examined three specific charge-changing perturbations, an evaluation of CI and AC approaches in λD would benefit from an additional future benchmark study incorporating a larger collection of charge-change perturbations, but which was beyond the scope of the present work. While the breadth of this study was limited to a small series of protein side chain perturbations, nonetheless, it provided valuable insight into the selection of a charge-correction strategy for λD simulations, creates an infrastructure for easy application of the CI correction in λD simulations, and defines a workflow for selecting which solvent molecules or ions are coupled to a primary charge-change perturbation. Together, these developments will be invaluable for further studies performing charge-change perturbations in λD. Our automated implementation of CI λD, along with input scripts necessary to reproduce this work, are available as an open source Zenodo repository. [66] The calculated ΔΔ*G*s for the EXOSC3 variants were rationalized by structural investigations of the λD simulation trajectories.

## METHODS

### Generating Exosome Structural Models

Initial coordinates for the RNA exosome complex model were obtained from a cryo-EM structure of the human nuclear exosome in complex with DIS3, MTR4, and MPP6 cofactors (PDB: 6D6Q). [16] The atomic coordinates of the exosome protein subunits were refined following a Rosetta macromolecular assembly refinement protocol developed by the DiMaio lab. [67,68] The MolProbity webserver was used for hydrogen atom placement as well as to identify suggested side chain flips for histidine, asparagine, and glutamine. [69–71] Manual inspection was performed to determine histidine protonation states (His-δ or His-ε) based on potential hydrogen bonding partners with consideration of the protonation states suggested by the Reduce program in MolProbity. [72] Propka was used to identify potential protonation state changes to Asp, Glu, Lys, Arg, and His residues. [73,74] Short protein breaks (≤ 6 amino acids) were repaired using Chimera’s Modeller tool. [75] RosettaES was used to rebuild a longer break in EXOSC3 (residues 39-60), [76] using a truncated exosome model spherically cut 40 Å away from EXOSC3. The resultant RosettaES-generated structure was then minimized and then further relaxed in a 900 ns MD simulation of the truncated exosome complex. EXOSC3 residues 30-70 were then incorporated back into the full exosome model and minimized. Residues 19-62 of the broken RNA strand were removed from the model and residues 19-59 were rebuilt using auto-DRRAFTER, [77,78] guided by the placement of the RNA strand present in the 6D6Q model as well as RNA from a yeast exosome structure (PDBID: 7AJU). [79] The rebuilt RNA was placed back into the full exosome model. Following the above changes, the system was relaxed by minimization followed by a 325 ns MD simulation. To decrease the computational cost of the exosome complex simulations, a spherically truncated form of the final exosome-bound model was created by discarding all residues outside a 27 Å radius from EXOSC3. The unbound EXOSC3 model was created by extracting EXOSC3 from the full model of the exosome complex. To measure ΔΔ*G*_fold_ with λ-dynamics, unfolded models of EXOSC3 were made by extracting perturbation site-centered pentapeptides from the full exosome model. [47] Specifically, the D132-, G135-, and G191-centered pentapeptides were formed by separately extracting residues 130-134, 133-137, or 189-193 from the unbound EXOSC3 model.

### System preparation

All systems were solvated with the TIP3P water model using the CHARMM-GUI solution builder tool. [80,81] Na+ and Cl-ions were added to neutralize the net charge of the system and achieve an ionic strength of 100 mM NaCl. A solvent buffer of 20 Å was used for the unfolded EXOSC3 (pentapeptide) models as CI simulations of charge-change perturbations within smaller systems (i.e. ligands) were found to have improved accuracy with larger box sizes. [55] All other models had a solvent buffer of 10 Å. Native N- and C-terminal residues were capped with standard CHARMM36 patches of GLYP, NTER, and CTER while ACE, ACP, and CT3 patches were used to cap protein and peptide chain termini in the truncated complex that did not coincide with a protein’s native N- and C-terminal residues. All proteins and nucleic-acids were modeled using the CHARMM36 all-atom force field. [82–84] All minimizations listed below were performed with the CHARMM molecular simulation package using the steepest descent algorithm. [62–64] Minimizations were performed over 1000 - 2000 steps split over stages, starting with restraints on all protein and nucleic acid atoms and then incrementally removing the harmonic restraints such that the final minimization was performed with restraints removed from all atoms. In the truncated exosome model, the terminal RNA bases and all protein backbone atoms greater than 17 Å away from EXOSC3 were harmonically restrained with a force constant of 10 kcal/ molÅ^2^ to maintain the natural shape and fold of the exosome complex surrounding EXOSC3. [62–64]

### Simulation details

All MD and λD simulations were performed with the CHARMM molecular software package [62–64] All simulations were performed with a timestep of 2 fs. Sampling in the isothermal isobaric (NPT) ensemble was performed with a Monte Carlo barostat and Langevin dynamics to maintain a pressure of 1.0 atm and temperature of 30 °C. [85,86] The SHAKE algorithm was employed to constrain all hydrogen-heavy atom bond lengths, and periodic boundary conditions were employed. [87] The particle mesh Ewald (PME) method was used to compute all long-range electrostatic interactions, and all nonbonded Lennard–Jones long-range interactions were truncated at 10 Å, with force-switching to zero between 9 and 10 Å. [59,88–90]

### λ-Dynamics Simulations

Alchemical perturbations were represented with hybrid multiple-topology models using the BLOCK facility within CHARMM, following standard procedures for λD simulations. [40–43,45–49,58,61,65,91] Full-residue perturbations were performed, such that all amino acid backbone atoms were included as alchemical substituent atoms, following a strategy developed for alchemical perturbations of proline and glycine residues with λD. [65] λD investigations of the EXOSC3 variants were separated by perturbation site. Within the D132 variant simulations, the native D132 was perturbed to an Ala residue. Within the G135 variant simulations, the native G135 was perturbed to α-aminobutyric acid (Abu), norleucine (Nle), and Arg residues. Abu and Nle served as intermediate states between Gly and Arg. Within the G191 variant simulations, the native G191 was perturbed to Ala, Cys, and Asp residues. Ala served as an intermediate state between Gly and the Cys and Asp residues. For calculating free-energy differences, λ states were saved every 10 steps and a cutoff value of λ ≥ 0.99 was used as an approximation of the physical end state λ = 1.0. Nonbonded interactions for the alchemical substituents were scaled by λ using a soft-core Lennard– Jones potential. [46] Substituent dihedral angles were scaled by λ, as this was previously found to yield better rotational sampling without causing conformations to become trapped in local energy minima. [46,49] Substituent bonds, angles, and improper dihedral angles were not scaled by λ. The Adaptive Landscape Flattening (ALF) algorithm was performed as follows to identify appropriate biasing potentials for each λ state. Optimized biases flatten the free energy landscape in λ-space and enables the dynamic sampling of many different perturbation end states. [46,47] Brief 100-ps pre-production simulations were performed for a minimum of 100 iterations or until a complete or nearly complete free energy landscape was produced for all λs by visual inspection. Pre-production simulations with a simulation length of 1 ns were then performed until the estimated free energy landscape was flattened to a range of ca. 1 kcal/mol with frequent sampling of the perturbation end states and the end state sampling ratio of the maximally and minimally sampled substituent for each perturbation site fell within a ratio of 10. This was then sequentially followed by five replicas of 5-ns, 25-ns, and, as needed, 100-ns pre-production λD simulations. Optimal biases were obtained with ALF after a cumulative of 170-800 ns of sampling (Table S2). Finally, λD production simulations were run for the exosome-bound, unbound, and unfolded EXOSC3 systems. The first fifth of each production simulation was excluded as equilibration prior to determination of the free energy changes. Final relative free energy differences were calculated with WHAM, [92] and statistical errors were estimated using bootstrapping.

### Analytic correction

A simplified post-simulation analytic correction (AC) was performed to account for the change in charge accompanying the D132A, G135R, and G191C perturbations. Following the completion of a routine λD simulation, a post-alchemical correction (Eq. 2) [52] was applied to the ΔΔ*G* results.

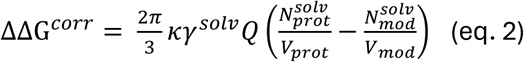

where 𝜅 is the electrostatic constant, 𝛾^solv^ is the quadrupole moment trace of the solvent model relative to a van der Waals interaction site (calculated as 0.764 e·Å^2^ for the TIP3P water model [59]), Q is the change in charge between alchemical end states. 𝑁^solv^ is the number of solvent molecules, and 𝑉 is the volume of the periodic box averaged over all production frames. “*prot”* denotes the exosome-bound EXOSC3 for the binding free energy simulations or the folded EXOSC3 state within the folding free energy simulations and “*mod”* denotes the unbound EXOSC3 state within the binding free energy simulations or an unfolded EXOSC3 pentapeptide within the folding free energy simulations.

### Co-alchemical ion

In the CI method, the substituents at a charge-change perturbation site are coupled to a perturbation of the opposite charge in the solvent [93] to maintain a neutral net charge of the system regardless of the identity of the side chain at the perturbation site. Charged residues D132, G135R, and G191 were each coupled to water molecules. Neutral D132, G191, G191C, and the neutral intermediate G191A were each coupled to a chloride atom. Neutral G135, G135Nle, and G135Abu were each coupled to a sodium atom. The non-hydrogen co-alchemical ion atom pairs, (Cl and O or Na and O), were restrained to one another using CHARMM’s constrained atoms (CATS) algorithm. [65] The alchemical solvent molecules were permitted to move around the simulation box freely.

### Structural analysis

Prior to all structural analyses of the exosome MD and λD simulations, frames within the first fifth of each simulation replica were discarded as equilibration. Production frames refer to the remaining four-fifth frames of each trajectory that were analyzed. Mean shift clustering [94] was used to quantify the frequency of the native L_23_ configuration in the D132 and D132A EXOSC3 variant frames. To summarize, the G134 Ψ, G135 Φ, and G135 Ψ dihedrals were collected from all frames. The Φ and Ψ dihedrals were scaled to range from 0 ° to 360 ° or from -60 ° to 300 °, respectively, to prevent the artificial separation of clusters due to the linearization of the cyclical dihedral measurement. Two rounds of mean shift clustering were then performed on the collected dihedrals with all datapoints assigned to clusters without discarding outliers. The datapoints for every cluster identified with the first round of mean shift clustering were then collected and re-analyzed by mean shift clustering with the same conditions as before. The native configuration cluster was identified by the cluster center nearest the G134 Ψ, G135 Φ, and G135 Ψ dihedral values of 20 °, 270 °, and 200 °, respectively. The analysis described above was separately performed for each D132 variant using an inhouse python script. MD and λD trajectories were analyzed with the assistance of PyMOL [95]. The 191 N-Y164 O, Y164 N-191 O, and I193 N-L162 O distances were collected from all G191, G191C, and G191D production frames. The Y109 N-G134 O and shortest R108 guanidine N-G134/G135 O atomic distances were collected from all production frames of the D132 EXOSC3 variant as well as the D132A production frames that mean shift clustering identified as having an alternative L_23_ configuration. All bar charts, violin plots, and scatter plots were generated using Matplotlib. [96]

## SUPPORTING INFORMATION

Figures tracking the change in ΔG over the last 50 ns of each λD simulation for the exosome-bound, unbound, and unfolded EXOSC3 variant models, D132A-induced alternative L_23_ conformational statistics, conformational changes in unbound G135R EXOSC3, distributions of the β_L5_-β_4_ hydrogen bonds for the unbound G191 variants, tables of the λD pre-production simulation times, calculated ΔΔ*G*s, and CI and AC λD simulation sampling statistics.

## AUTHOR CONTRIBUTIONS

M.P.B. – experimental design, data acquisition, data analysis, writing and revision of manuscript

H.R.S.W. - experimental design, data acquisition, data analysis

A.M.R. – experimental design, data acquisition, data analysis, writing and revision of manuscript

K.M.C. – data analysis, writing and revision of manuscript

A.L.M. – funding acquisition, project design, experimental design, data analysis, writing and revision of manuscript

J.Z.V. – funding acquisition, project design, experimental design, data analysis, revision of manuscript

## ACKNOWLEDGEMENTS

The authors gratefully acknowledge the National Institutes of Health for support of this work, through several funding mechanisms reported above. The authors acknowledge the Indiana University Pervasive Technology Institute for providing supercomputing and storage resources that have contributed to the research results reported within this paper. The mass spectrometry work was performed by the Indiana University School of Medicine Center for Proteome Analysis.

Acquisition of IUSM Center for Proteome Analysis’ instrumentation was provided by the Indiana University Precision Health Initiative and the Indiana Clinical Translational Sciences Institute. The Center for Proteome Analysis also receives support from 3UL1TR002529 and P30CA082709. The authors acknowledge and thank Whitney R. Smith-Kinnaman and Emma H. Doud for their assistance with paired proteomics work and Ryan L Hayes for a discussion about the simplified AC correction scheme.

