## Supporting Information for "Quantifying the Structural and Energetic Consequences of EXOSC3 S1 Domain Variants from a Comparative Assessment of λ-Dynamics with Two Charge-Changing Perturbation Strategies"

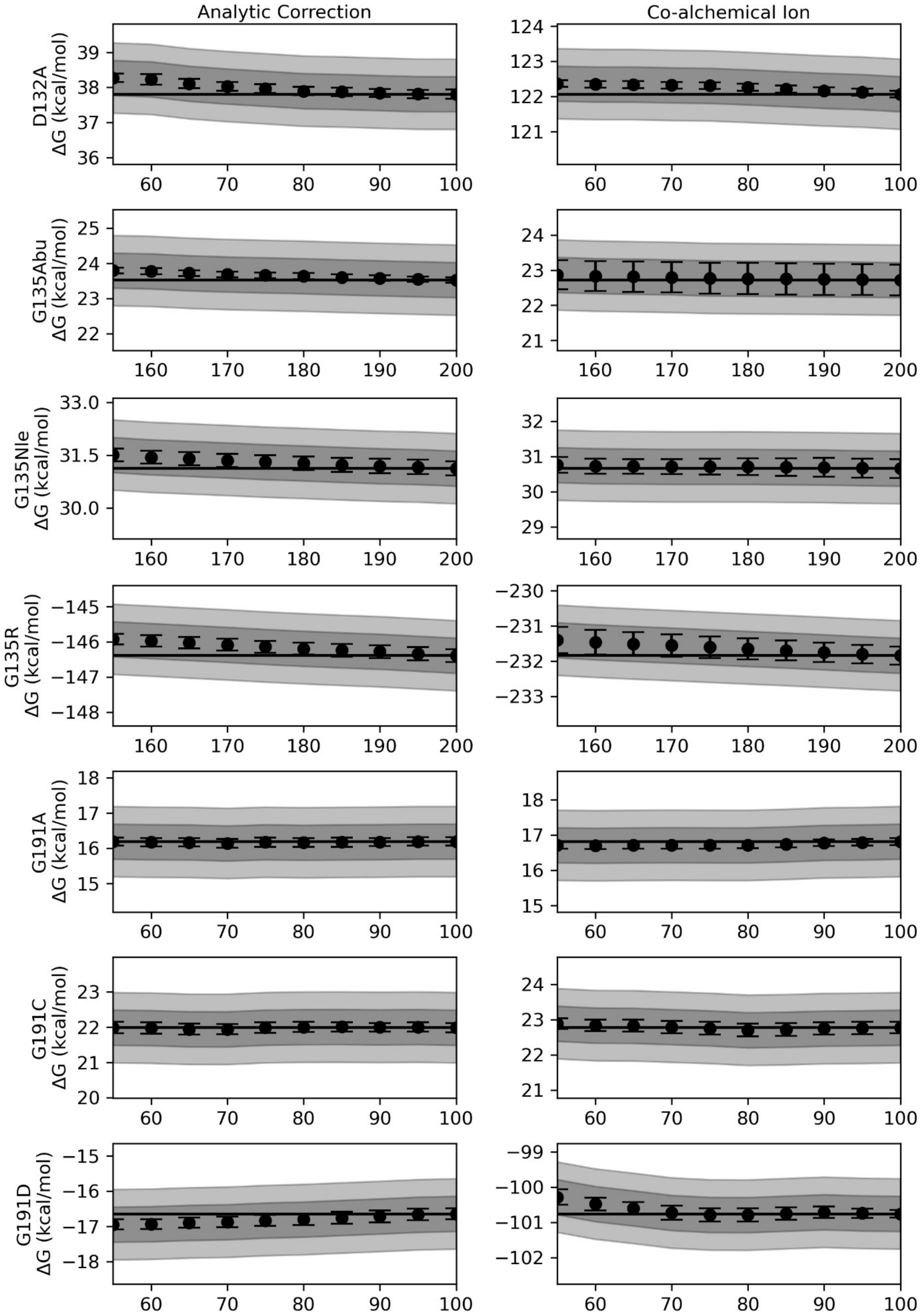
Analytic Correction
Co-alchemical Ion
D132A
ΔG (kcal/mol)
G135Abu
ΔG (kcal/mol)
G135Nle
ΔG (kcal/mol)
G135R
ΔG (kcal/mol)
G191A
ΔG (kcal/mol)
G191C
ΔG (kcal/mol)
G191D
ΔG (kcal/mol)

**Figure S1: Convergence plots for exosome-bound EXOSC3 calculations.** Changes in free energy differences ($\Delta G$) computed with λ-dynamics and analytic correction (left column) or co-alchemical ion (right column) charge-change correction schemes are graphed as a function of time for all exosome-bound EXOSC3 variants. The solid horizontal line marks the final $\Delta G$ value. Dark and light gray bands represent $\Delta G$ deviations within ± 0.5 and ± 1.0 kcal/mol of calculated free energy values, respectively. All plots cover the last 50 nanoseconds of each simulation with a y-axis scale fixed at 4 kcal/mol centered on the final computed $\Delta G$ value.

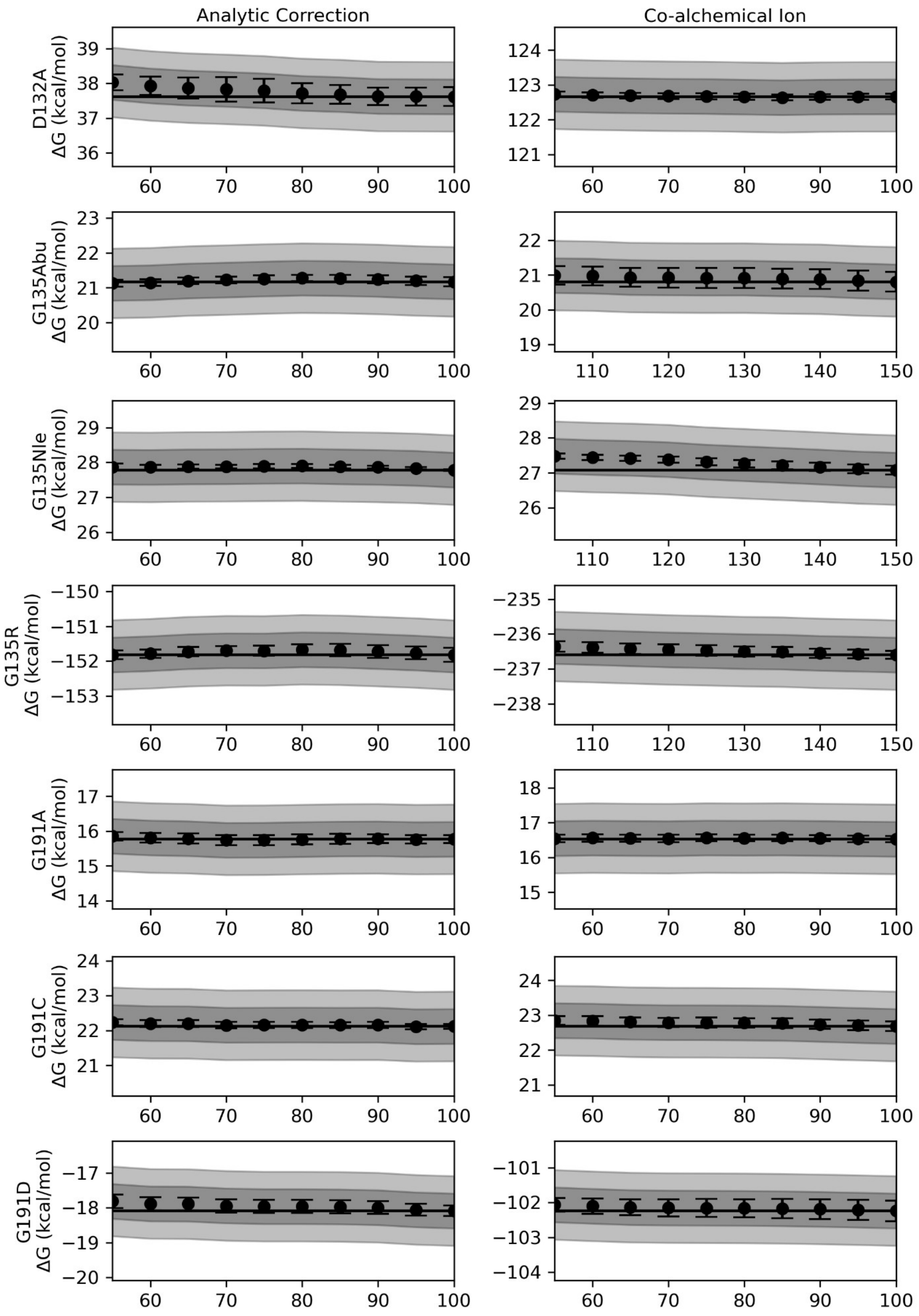

Analytic Correction
Co-alchemical Ion
D132A
ΔG (kcal/mol)
G135Abu
ΔG (kcal/mol)
G135Nle
ΔG (kcal/mol)
G135R
ΔG (kcal/mol)
G191A
ΔG (kcal/mol)
G191C
ΔG (kcal/mol)
G191D
ΔG (kcal/mol)

**Figure S2: Convergence plots for unbound EXOSC3 calculations.** Changes in free energy differences (Δ*G*) computed with λ-dynamics and analytic correction (left column) or co-alchemical ion (right column) charge-change correction schemes are graphed as a function of time for all unbound EXOSC3 variants. The solid horizontal line marks the final Δ*G* value. Dark and light gray bands represent Δ*G* deviations within ± 0.5 and ± 1.0 kcal/mol of calculated free energy values, respectively. All plots cover the last 50 nanoseconds of each simulation with a y-axis scale fixed at 4 kcal/mol centered on the final computed Δ*G* value.

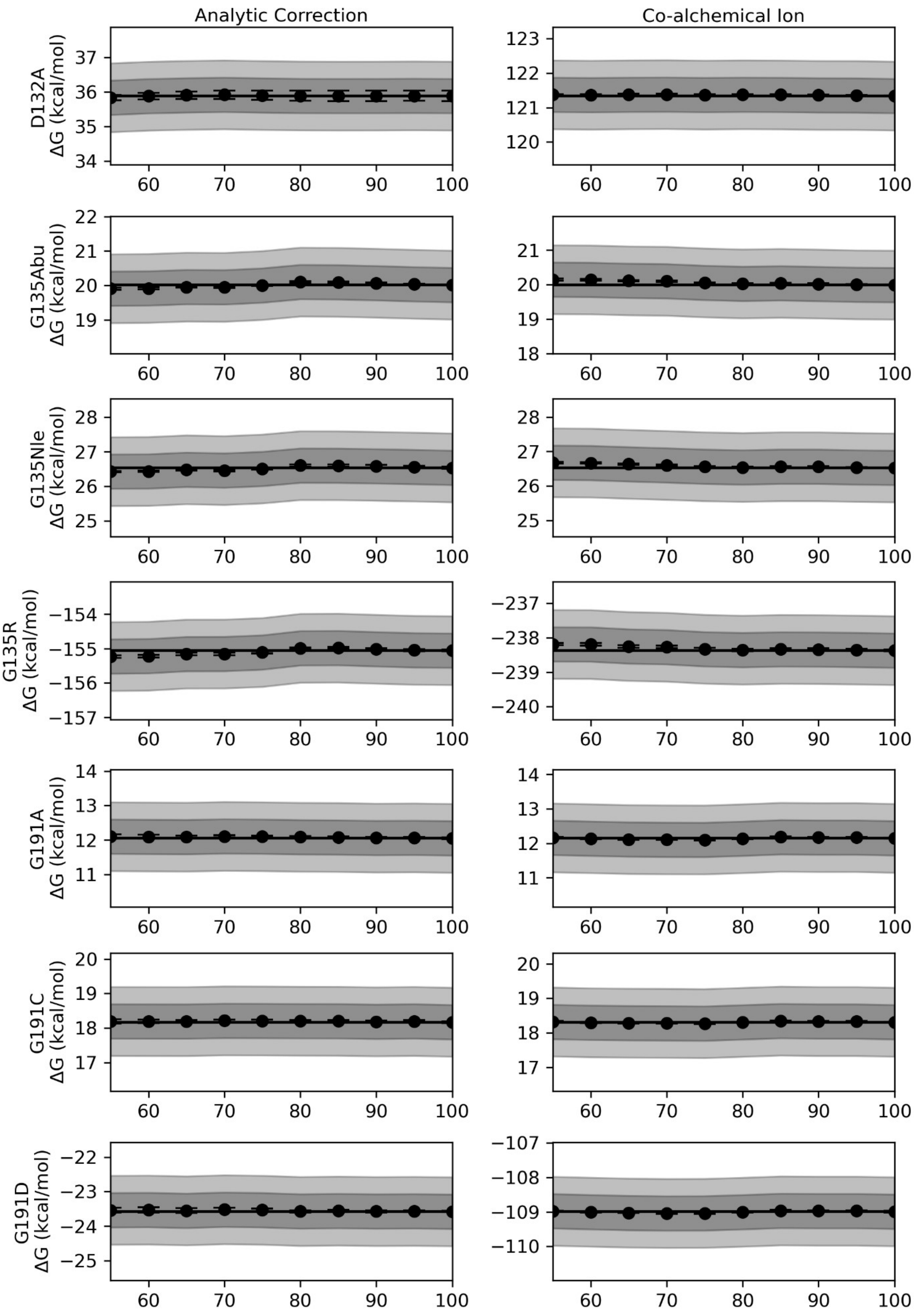
Analytic Correction
Co-alchemical Ion
D132A ΔG (kcal/mol)
G135Abu ΔG (kcal/mol)
G135Nle ΔG (kcal/mol)
G135R ΔG (kcal/mol)
G191A ΔG (kcal/mol)
G191C ΔG (kcal/mol)
G191D ΔG (kcal/mol)
60
70
80
90
100

**Figure S3: Convergence plots for unfolded EXOSC3 calculations.** Changes in free energy differences ($\Delta G$) computed with λ-dynamics and analytic correction (left column) or co-alchemical ion (right column) charge-change correction schemes are graphed as a function of time for all unfolded EXOSC3 variants. The solid horizontal line marks the final $\Delta G$ value. Dark and light gray bands represent $\Delta G$ deviations within ± 0.5 and ± 1.0 kcal/mol of calculated free energy values, respectively. All plots cover the last 50 nanoseconds of each simulation with a y-axis scale fixed at 4 kcal/mol centered on the final computed $\Delta G$ value.

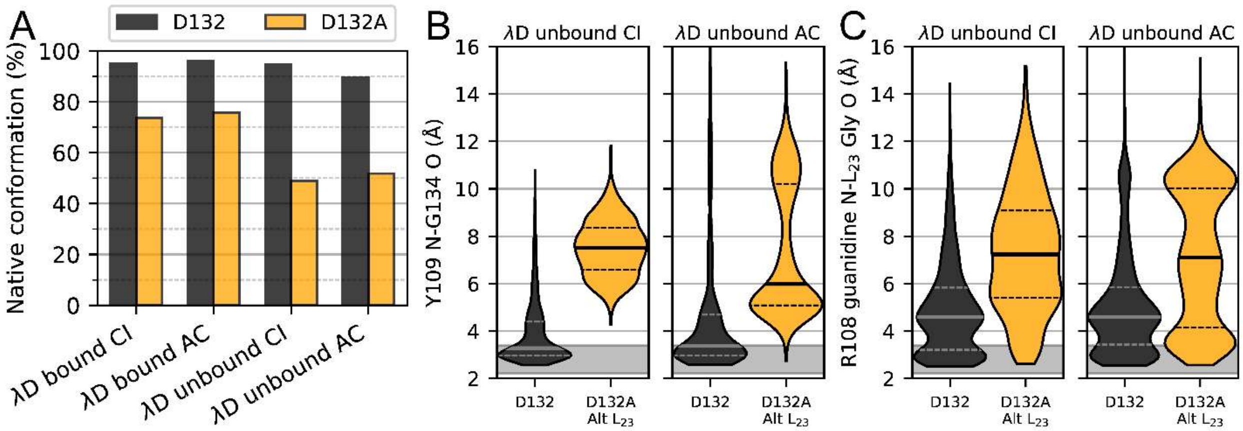


**Figure S4: D132A-induced alternative $L_{23}$ conformation statistics**. (A) Quantification of the frequency of the native $L_{23}$ configuration in D132 (black bars) and D132A (orange bars) frames in all λD simulations of exosome-bound and unbound EXOSC3. (B and C) Distributions of distances between the NT/S1 linker and $L_{23}$ hydrogen bonding partners for CI- and AC-corrected λD simulations of unbound EXOSC3 D132 and D132A variants. (B) Distance distributions for the G134 O-Y109 N hydrogen bond. (C) Distance distributions of the shortest distance between the R108 guanidine nitrogens and the $L_{23}$ G134 or G135 oxygens. All violin plots are normalized to have the same width; solid lines represent median values, and dashed lines show the interquartile range. The gray band highlights a typical hydrogen bond distance between 2.2 and 3.4 Å heavy atom separation. Measurements were made for all D132 frames (D132) and all D132A frames with an alternative $L_{23}$ configuration (D132A Alt $L_{23}$).

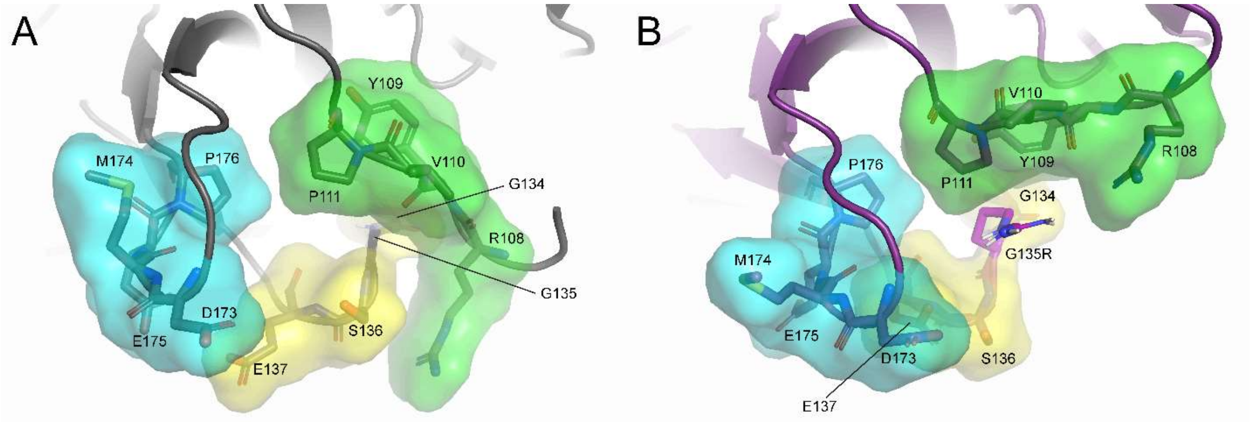


**Figure S5. Disruption of packing interactions in unbound EXOSC3 by the G135R mutation**. Representative unbound EXOSC3 frames show packing interactions between $L_{23}$, $L_{45}$, and the NT/S1 linker in native EXOSC3 (A) and G135R EXOSC3 (B). Exosome complex proteins are colored light gray, G135 EXOSC3 is colored dark gray, and G135R EXOSC3 is colored purple. Interacting residues within the $L_{23}$, $L_{45}$, and the NT/S1 linker are labeled and represented as sticks. A color-coded surface is shown: NT/S1 linker (green), $L_{23}$ (yellow), and $L_{45}$ (cyan).

**Table S1: λ-Dynamics Computed Relative Free Energies of Binding ($\Delta\Delta G_{bind}$) and Folding ($\Delta\Delta G_{fold}$) for EXOSC3 S1 Domain variants with CI and AC Charge-Change Correction Schemes**

| | $\Delta\Delta G_{bind} \pm \sigma$ (kcal/mol) | | $\Delta\Delta G_{fold} \pm \sigma$ (kcal/mol) | |
|---|---|---|---|---|
| EXOSC3 variant | CI | AC | CI | AC |
| D132 | 0.00 ± 0.265 | 0.00 ± 0.274 | 0.00 ± 0.308 | 0.00 ± 0.116 |
| D132A | 0.19 ± 0.297 | 0.07 ± 0.129 | 1.73 ± 0.308 | 1.97 ± 0.085 |
| G135 | 0.00 ± 0.356 | 0.00 ± 1.022 | 0.00 ± 0.291 | 0.00 ± 0.992 |
| G135Abu | 2.35 ± 0.150 | 1.91 ± 0.525 | 1.16 ± 0.128 | 0.82 ± 0.284 |
| G135Nle | 3.34 ± 0.199 | 3.58 ± 0.295 | 1.25 ± 0.038 | 0.55 ± 0.126 |
| G135R | 5.43 ± 0.272 | 5.43 ± 0.277 | 3.23 ± 0.202 | 2.40 ± 0.118 |
| G191 | 0.00 ± 0.231 | 0.00 ± 0.328 | 0.00 ± 0.121 | 0.00 ± 0.131 |
| G191A | 0.43 ± 0.164 | 0.29 ± 0.133 | 3.71 ± 0.116 | 4.38 ± 0.088 |
| G191C | -0.13 ± 0.147 | 0.09 ± 0.214 | 3.95 ± 0.074 | 4.37 ± 0.138 |
| G191D | 1.45± 0.221 | 0.81 ± 0.311 | 5.49 ± 0.153 | 6.12 ± 0.294 |

**Table S2: Analysis of Total ALF Pre-Production Simulation Times for CI and AC λD Simulations**

| Perturbation set | Total CI pre-production simulation time (ns) | Total AC pre-production simulation time (ns) |
|---|---|---|
| Model | Exosome-bound EXOSC3 | |
| D132 | 195.2 | 228.0 |
| G135 | 772.0 | 754.4 |
| G191 | 264.4 | 478.0 |
| Average | 410.5 | 486.8 |
| Model | Unbound EXOSC3 | |
| D132 | 217.2 | 172.5 |
| G135 | 790.0 | 226.0 |
| G191 | 795.0 | 298.0 |
| Average | 600.7 | 232.2 |
| Model | Mutation site-centered EXOSC3 pentapeptide | |
| D132 | 169.4 | 260.4 |
| G135 | 301.0 | 292.0 |
| G191 | 276.0 | 301.0 |
| Average | 248.8 | 284.5 |
| Total Average | 420.0 | 334.5 |

**Table S3: Analysis of CI and AC λD Production Simulation Sampling Statistics**

| | CI | | AC | |
|---|---|---|---|---|
| EXOSC3 variant | Sampling (% simulation) | # Transitions per 100 ns | Sampling (% simulation) | # Transitions per 100 ns |
| Model | Exosome-bound EXOSC3 | | | |
| D132 | 26.9 | 45.0 | 6.1 | 93.8 |
| D132A | 19.5 | 45.0 | 29.9 | 94.2 |
| G135 | 6.1 | 6.1 | 16.2 | 26.9 |
| G135Abu | 0.2 | 36.4 | 0.6 | 58.2 |
| G135Nle | 0.3 | 44.8 | 1.0 | 59.4 |
| G135R | 10.2 | 25.4 | 14.0 | 24.9 |
| G191 | 5.0 | 379.0 | 4.1 | 494.0 |
| G191A | 3.1 | 1072.6 | 4.8 | 1243.2 |
| G191C | 2.5 | 920.2 | 3.7 | 977.6 |
| G191D | 3.4 | 36.6 | 2.0 | 78.2 |
| Average | 7.7 | 261.1 | 8.2 | 315.0 |
| Model | Unbound EXOSC3 | | | |
| D132 | 9.1 | 75.8 | 18.3 | 260.4 |
| D132A | 10.6 | 76.4 | 19.2 | 260.0 |
| G135 | 5.2 | 186.2 | 1.5 | 57.2 |
| G135Abu | 3.5 | 376.2 | 1.9 | 295.6 |
| G135Nle | 2.0 | 284.8 | 3.9 | 408.4 |
| G135R | 3.1 | 62.4 | 13.3 | 233.0 |
| G191 | 3.3 | 337.8 | 4.3 | 602.2 |
| G191A | 3.1 | 1100.0 | 2.5 | 1471.2 |
| G191C | 2.6 | 1032.8 | 3.9 | 1276.2 |
| G191D | 2.9 | 112.0 | 5.4 | 137.2 |
| Average | 4.5 | 364.4 | 7.4 | 500.1 |
| Model | Mutation site-centered EXOSC3 pentapeptide | | | |
| D132 | 5.1 | 100.0 | 13.7 | 625.8 |
| D132A | 8.3 | 100.0 | 22.9 | 625.8 |
| G135 | 3.5 | 359.6 | 3.0 | 278.4 |
| G135Abu | 3.4 | 719.6 | 3.6 | 781.0 |
| G135Nle | 3.7 | 694.4 | 3.8 | 815.8 |
| G135R | 2.1 | 151.6 | 3.8 | 503.8 |
| G191 | 2.3 | 551.4 | 4.3 | 1221.6 |
| G191A | 2.7 | 1283.2 | 3.9 | 2145.4 |
| G191C | 3.4 | 1265.4 | 3.9 | 2008.4 |
| G191D | 2.5 | 123.6 | 3.4 | 507.2 |
| Average | 3.7 | 534.9 | 6.6 | 951.3 |
| Total Average | 5.3 | 386.8 | 7.4 | 588.8 |
